# Glycogen phosphorylase from the methanogenic archaeon *Methanococcus maripaludis*: Unique regulatory properties of a pyridoxal 5’-phosphate independent phosphorylase

**DOI:** 10.1101/2024.09.29.615707

**Authors:** Felipe González-Ordenes, Nicolas Herrera-Soto, Leslie Hernández-Cabello, Catalina Bustamante, Gabriel Vallejos-Baccelliere, Victor Castro-Fernandez, Victoria Guixé

**Affiliations:** Laboratorio de Bioquímica y Biología Molecular, Departamento de Biología, Facultad de Ciencias, Universidad de Chile

**Keywords:** Glycogen phosphorylase, pyridoxal 5’-phosphate, *Methanococcus maripaludis*, enzyme regulation, glycogen, archaea

## Abstract

Glycogen phosphorylase (GP) is a critical enzyme in glycogen metabolism. Even though methanogens from the archaeal orders *Methanosarcinales* and *Methanococcales* are unable to grow on sugars, they store glycogen, which is metabolized through the glycogenolysis and glycolytic pathways when the carbon source for methanogenesis is depleted. Under these metabolic conditions, the activity of the GP enzyme is essential. To be active, all phosphorylases characterized to date require the cofactor pyridoxal 5’-phosphate (PLP). This cofactor is covalently bound via Schiff base to a strictly conserved lysine residue at the active site. Extensive GP sequence analysis of organisms from different domains of life shows strict conservation of active site residues despite significant differences in sequence length. Interestingly, in GP sequences of organisms from the order *Methanococcales* of archaea, a threonine residue replaces the conserved lysine involved in PLP binding. The purification and characterization of recombinant GP from *Methanococcus maripaludis* show that the enzyme exhibits glycogen phosphorylase activity and high specificity for glycogen as a substrate. Analysis of the PLP content performed by several methods, such as absorbance, fluorescence, cyanohydrin adduct formation, and mass spectrometry, confirmed the absence of PLP. The results demonstrate that an archaeal GP from the order *Methanococcales* performs catalysis without the PLP cofactor, deviating from the well-established phosphorylase catalytic mechanism and revealing new scenarios for the glucosyltransferase reaction. Moreover, analysis of enzyme regulation shows that the activity is affected by various molecules, including nucleotides, intermediates of carbon metabolism, and phosphate species. Most of these molecules have not previously been identified as regulators of glycogen phosphorylases in prokaryotes. These results suggest that other GPs from *Methanococcales* can undergo complex regulation.

## 1. Introduction

Glycogen is a branched glucose polymer that constitutes a carbon reservoir widespread in eukaryotes, bacteria, and archaea. Under nutrient starvation conditions, glycogen allows prokaryotes to generate energy and synthesize building blocks. In humans, glycogen provides muscle contraction energy, maintains glucose homeostasis [1], and supports cognitive processes such as learning and memory [2].

The first step in glycogen degradation is performed by Glycogen Phosphorylase (GP) (EC 2.4.1.1), which catalyzes the rate-limiting step of glycogenolysis, releasing glucose-1-phosphate from the terminal alpha-1,4-glycosidic bond. All phosphorylases characterized to date require the cofactor pyridoxal 5’-phosphate (PLP) to perform catalysis, which is covalently bound to an active site conserved lysine residue via a Schiff base [3].

The role of PLP in phosphorylase catalysis differs from other PLP-dependent enzymes, which are based on the reactivity of the aldehyde group of PLP [4]. Seminal work on GPs demonstrated that reduction of the imine between the 4’-aldehyde and the ε-amino group of lysine resulted in no substantial loss of activity [5]. At the same time, the reconstitution of GP with different chemical analogs of PLP modified in every position around the pyridine ring identified the 5’-phosphate group as essential for catalysis [5,6]. According to the proposed catalytic mechanism for the phosphorylase reaction, the phosphate group of PLP acts as a general acid and protonates the substrate phosphate, operating as a proton shuttle [6,7]. Besides its catalytic role in phosphorylases, it has also been reported that PLP fulfills a role in stabilizing the quaternary structure [5,8].

Due to its role in glucose homeostasis, vertebrate phosphorylases are tightly regulated by phosphorylation and multiple allosteric effectors, like AMP, ATP and glucose 6-phosphate. X-ray crystallography has revealed four regulatory sites in GP from rabbit muscle, the model isoenzyme. In contrast, evidence of regulation in other phylogenetic groups points to a simpler regulatory strategy. In more primitive eukaryotes, such as yeast, GP is also regulated by phosphorylation [9] and a small set of effectors that includes glucose-6-phosphate and UDP-glucose [10,11]. In bacteria, regulation has been demonstrated in a small subset of enzymes that respond to ADP-glucose and UDP-glucose and exhibit a slight effect by AMP [12,13]. For archaeal enzymes, allosteric regulation has not been evaluated, although glycogen accumulation has been described in several phylogenetic groups [14,15], including autotrophic organisms that do not assimilate carbohydrates, like methanogens [16]. Whole-genome sequencing of these organisms revealed that, even when they do not assimilate carbohydrates, all enzymes required for glycolysis/gluconeogenesis are present, including genes for glycogen synthesis and degradation, as evidenced by species like *Methanosarcina acetivorans* (order *Methanosarcinales*) and *M. maripaludis* (order *Methanococcales*). Microbiological studies of methanogens have demonstrated glycogen synthesis under carbon excess and nitrogen limitation conditions. However, upon depletion of the carbon source employed for methanogenesis, glycogen consumption is initiated concomitantly with an increase in the glycolytic flux and the reactivation of methanogenesis [15,17], a process for which the activity of the GP enzyme is essential. Considering the importance of glycogen in the survival and adaptation of methanogenic organisms, the need for tight regulation of glycogen metabolism, it is essential to address the kinetic and regulatory properties of glycogen phosphorylases from these organisms.

A multiple sequence alignment of GPs from eukaryotes, bacteria, and archaea revealed a strict conservation of active site residues across all phosphorylase sequences. Noticeably, in enzymes belonging to organisms from the order *Methanococcales* of archaea, a threonine residue replaces the strictly conserved lysine involved in PLP binding, raising questions about the ability of glycogen phosphorylases from *Methanococcales* to bind the cofactor and perform catalysis in its absence.

Characterization of the recombinant glycogen phosphorylase from *Methanococcus maripaludis* (MmGP) shows that MmGP generates glucose-1-phosphate, prefers large glucose polymers like glycogen, and has a specific activity similar to GPs from archaea and bacteria. However, absorbance, fluorescence, and mass spectrometry assays indicate that the enzyme lacks the PLP cofactor. Screening for potential effectors identified six inhibitory metabolites and one activator. These findings reveal unique properties of a phosphorylase from the *Methanococcales* order, capable of catalysis without the cofactor pyridoxal 5’-phosphate, along with unique regulatory properties.

## 2. Material and methods

### Materials

Unless otherwise specified, all materials were from Sigma Aldrich (Merck KGaA, Darmstadt, Deutschland) at the highest available purity. Maltotetraose, pentaose and hexaose were from Aaron Chemicals (San Diego, USA). Glucose-6-phosphate dehydrogenase from *Leuconostoc mesenteroides* (*Lm*G6PDH) was recombinantly expressed in *E. coli* BL21 cells and purified through Ni-NTA affinity chromatography in our laboratory.

### Sequence alignment, SeqLogo analysis and phylogenetic inference

Phosphorylase sequences were collected from the non-redundant NCBI protein database using the search algorithm in November 2021 [18]. The analysis was conducted with the aim of enriching the search with archaeal sequences. Consecutive searches were carried out on the main phylogenetic orders of the *Thermoproteota* and *Euryarchaeota* groups, while for eukaryotes and bacteria, the sequence search was limited to those representative groups already studied in the literature, resulting in 94 final sequences. The alignment was performed using the Multiseq program [19] implemented in VMD using the ClustalW algorithm [20]. The sequence logo analysis was created using the “Weblogo” server [21], while the phylogenetic tree for the 94 aligned sequences was inferred by the maximum likelihood method implemented in PhyML v 3.3 [22] using as evolutionary model the LG matrix [23] with + I + G and F parameters.

### Protein expression and purification

The gene for MmGP (Uniprot code Q6LXX5) was synthesized by GenScript USA Inc. (Piscataway, NJ, United States), codon-optimized for expression in *Escherichia coli*, and subsequently cloned into the vector pET-TEV-28a(+) using the restriction sites NdeI and BamHI) to incorporate a His-tag and a TEV protease cleavage-site. Chemically competent cells of *E. coli* BL21(DE3) were transformed with the expression vector and subsequently grown overnight at 37°C for 16 h in 1L of Luria–Bertani broth (LB), supplemented with 30 µg/ml of kanamycin. Then, protein expression was performed for 4 h at 37°C and was initiated with 1 mM IPTG. The cells were centrifuged for 10 min at 4000 g, and the pellet was resuspended in 50 ml of buffer A composed of 50 mM glycine pH 9.5, 500 mM NaCl, and 20 mM imidazole and cells were disrupted by sonication. The lysate was centrifuged at 18000 g for 30 min, and the supernatant was filtered through a 0.45 µm membrane and loaded onto a nickel affinity column (5 mL GE Healthcare HisTrap). The protein was eluted using a gradient from 20 to 500 mM imidazole (solution B: 500 mM NaCl, 500 mM imidazole and 25 mM Tris-HCl pH 7.8). Fractions with phosphorylase activity were collected according to the analysis of their purity by SDS-PAGE. Protein concentration was determined by the Bradford assay using Bovine Serum Albumin (BSA) as standard.

### Size exclusion chromatography

The Hi-Prep 16/60 Sephacryl S-200 column (GE-Healthcare) was equilibrated with buffer 25 mM glycine, 300 mM NaCl and at pH 9.0. The column was calibrated using a commercial mixture of molecular markers: horse myoglobin, chicken ovalbumin, and bovine gamma globulin (BioRad Gel Filtration Standard #1511901). Elution volumes were determined for each marker, and the molecular weight of MmGP was calculated by interpolating in the linear regression of the calibration curve (log (PM) versus retention volume).

### Glycogen phosphorylase activity and kinetic characterization

Glycogen phosphorylase activity was measured in a continuous coupled assay in the direction of glycogen breakdown by the photometrical determination of the rate of NADH formation at 340 nm. The enzymes of the coupled assay consisted of phosphoglucomutase (PGM from rabbit muscle, Sigma-Aldrich P3397) and glucose-6-phosphate dehydrogenase (G6PDH from *L. mesenteroides*). The standard assay conditions contained 50 mM imidazole buffer pH 7.0, 5 mM KH_2_PO_4_, 0.5 mM NAD+, 5 U G6PDH, 10 U PGM, and 5 mg/mL glycogen from oyster. Initial rates were determined from the increase in NADH absorbance using an extinction coefficient of 6.22 mM^-1^ cm^-1^ and corrected by the path length. Measurements were performed in an Agilent 8454 diode array spectrophotometer (Agilent, Santa Clara, CA, USA) at 25 °C, and one enzyme unit (U) was defined as the production of 1 µmol of product per minute. Glucose polymer specificity was evaluated by using molecules of different chain lengths and branching frequency at a fixed concentration of 1.5 mg/mL (oyster glycogen, bovine glycogen, amylopectin, and maltodextrins) or at 3.0 mg/mL (maltotriose, maltotetraose and maltopentaose) in standard assay conditions. The results were normalized considering 100% of the specific activity obtained with oyster glycogen. The optimum pH was determined using a Universal Buffer composed of Imidazole 100 mM, Tris-HCl 100 mM, Bis-Tris-HCl 100 mM, and sodium acetate 100 mM in standard assay conditions. A mixture of imidazole, Tris, Bis-Tris, and concentrated sodium acetate was prepared in a total volume of 30 mL. The solution was adjusted using a pH meter while HCl was gradually added to lower the pH. Aliquots were collected at 0.5 pH unit intervals, and their volumes were adjusted to achieve a final concentration of 1.0 M for each of the four components.

Kinetic parameters were determined by initial velocity measurements at different concentrations of KH_2_PO_4_ or glycogen at a saturating co-substrate concentration. The Michaelis-Menten (equation 1) or the substrate inhibition equation (equation 2) was fitted to the data by nonlinear regression. The data was analyzed using Excel or GraphPad Prism 9 (Dotmatics Inc, Boston,MA, USA).

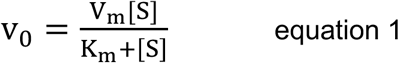

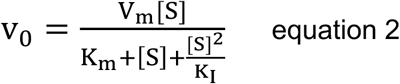

Effector screening was performed by measuring enzyme activity at 2 mM of each potential effector, using 0.3 mM KH2PO4 and 0,3 mg/mL glycogen. Dose-response curves were perfomed at 0.3 and 5 mM KH2PO4, with a fixed glycogen concentration of 0.3 mg/mL. IC50 (the concentration of effector at which half of the inhibitory effect is reached) and AC50 (the effector concentration at which half of the activator effect is reached) were determined by non-linear regression analysis. For curves showing only inhibition, either a hyperbolic decay with an asymptote into zero (equation 3) or one with an asymptote different from zero (equation 4) was fitted, where [L] correspond to the concentration of potential effector, v_0_^0^ corresponds to the initial velocity in the absence of effector, and v_0_^∞^ corresponds to enzyme activity extrapolated at infinite effector concentration. The best fit was selected using the AIC information criteria.

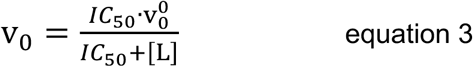

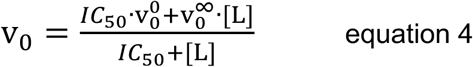

For activation-inhibition effect of ligands, the phenomenological equation (equation 5) involving both behaviors were fitted.

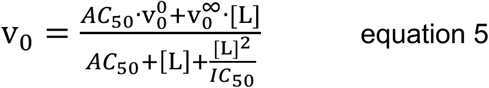

### Intrinsic and Extrinsic Fluorescence Analysis of MmGP

Fluorescence measurements of glycogen phosphorylase from *M. maripaludis* and glycogen phosphorylase *b* from rabbit muscle (*Oc*GP) were performed using 10 µM protein concentration in a JASCO J8300 spectrofluorometer. Protein concentrations were estimated using absorbance at 280nm (A_280_). Theoretical molar extinction coefficients were estimated from the protein sequence (ε=119640 M^-1^cm^-1^ for OcGP and 111730 M^-1^cm^-1^ for MmGP). For the measurements, *Mm*GP was dialyzed twice against 50 mM sodium phosphate pH 7.0 and 100 mM NaCl. OcGP was prepared from lyophilized powder (Sigma-Aldrich P6635) in buffer 50 mM sodium phosphate pH 7.0, 300 mM NaCl, 1 mM EDTA, and 0.5 mM 2-Mercaptoethanol. The fluorescence spectra of each buffer (blank) were subtracted from each sample using Spectramanager software (JASCO Inc., Japan). Tryptophan/tyrosine fluorescence was recorded at an excitation wavelength of 290 nm, while PLP fluorescence was collected using 290, 330 and 420 nm of excitation wavelengths.

### Quantification of the pyridoxal phosphate content by cyanohydrin formation

The assay of cyanohydrin formation was conducted as described previously [24,25]. Briefly, protein samples and PLP standards were prepared in 200 µL of nanopure water, then 200 µL of 11% trichloroacetic acid was added and incubated at 50°C for 15 minutes to denature the protein and release the PLP. The solution was neutralized to pH 7.5 with 140 µL of 3.3 M K_2_HPO_4_ and 3 µL of 1 M HEPES pH 7,8. The cyanohydrin formation was initiated with 50 µL of 20 mM KCN and incubated at 50 °C for 25 minutes. Then, pH was adjusted to 3.8 to enhance the fluorescence yield of the adduct by adding 70 µL of 28% v/v H_3_PO_4_ and 1 mL of 2.0 M potassium acetate-acetic acid buffer pH 3.8. Finally, the samples were centrifuged at 12,000g for 20 minutes, and the supernatant was stored at −20°C for analysis.

The quantification of adduct formation was performed by collecting the fluorescence emission spectra using a JASCO 8300 spectrofluorometer, with excitation at 325 nm and emission between 350 and 600 nm. To quantify the PLP content in enzyme samples, a calibration curve with PLP standards was built, from 0.1 to 1.0 nanomole in 700 µL, using the fluorescence at 420 nm.

### Determination of PLP content by mass spectrometry analysis

For the mass spectrometry (MS) analyses, 1.1 mg/mL of MmGP in 50 mM glycine pH 9.0, 300 mM NaCl, and 5% glycerol was sent for analysis to BGI Americas Corp (Cambridge, MA, USA). The whole-protein mass analysis was performed using the UHPLC Flex Vanquish system (ThermoFisher) coupled to an Orbitrap Q Exactive HF-X mass spectrometer (Thermo Fisher Scientific). Reverse phase chromatography was carried out on a MabPac column, with a flow rate of 300 µL/min and an injection volume of 3 µL. The following buffers were used:

Buffer A (H₂O with 0.1% formic acid) and Buffer B (ACN with 0.1% formic acid). The elution gradient was set to vary from 20% to 80% of Buffer B over 10 minutes, with the column maintained at 80°C and monitoring at 215, 254, and 280 nm. Sample ionization was performed using electrospray ionization (ESI) at a voltage of 3900 V and a capillary temperature of 250°C. The Orbitrap mass spectrometer was configured to operate in positive ion mode using a scan range of 800-6,000 m/z, with a combination of 10 microscans, achieving a resolution of 15,000. Data was acquired using Thermo Xcalibur Qual Browser software, and mass spectra deconvolution was performed using Thermo BioPharma Finder 4.0 with the ReSpect algorithm. As a quality control measure, 500 ng of IgG (Waters, 186006552) was analyzed.

For peptide analysis, the sample was denatured with 5% SDS and 100 mM TEAB buffer, reduced, alkylated, and then enzymatically digested overnight with a mixture of trypsin and Lys-C proteases. The digested fragments were subsequently separated using an S-Trap microcentrifuge column (Protifi, USA). Peptide separation was performed using an Ultimate 3000 UHPLC system (Thermo Fisher Scientific) with an Aurora C18 column (1.7 µm, 25 cm x 75 µm). The following mobile phases were used: Buffer A (2% acetonitrile with 0.1% formic acid) and Buffer B (80% acetonitrile with 0.1% formic acid). The elution gradient was programmed as follows: 2% to 35% Buffer B over 100 minutes, 35% to 65% Buffer B from 100 to 110 minutes, and 100% Buffer B from 110 to 120 minutes, with a flow rate of 0.3 µL/min. Ionization was performed using ESI, with the Orbitrap mass spectrometer operating in positive ion mode, full MS resolution at 120,000 and a mass range of 350-1,500 m/z. MS2 spectra were acquired at a resolution of 30,000 and fragmented using a fixed higher-energy collisional dissociation (HCD) collision energy of 25%, with a threshold of 2.5×10^4^ and considering charge states from 2 to 6. Peptides were analyzed to identify chemical modifications and sequence coverage using PMI Byos software (Protein Metrics). Raw data were searched against the sequence, accounting for general post-translational modifications (oxidation and deamidation) and expected modifications for PLP. Identified peptides were validated using the retention times of technical replicates, as well as full MS and MS2 data. Instrument quality control was conducted with 100 ng of HeLa standard.

### Molecular modelling

APO model for MmGP (UNIPROT Q6LXX5) was downloaded from the AlphaFold database (AF-Q6LXX5-F1-v4) [26] and was structurally superimposed with *Escherichia coli* maltodextrin phosphorylase (EcMalP, PDB 1L6I) [27] using pymol v2.07. PLP phosphoryl, phosphate, maltopentose (M5) and crystallographic waters of the active site of 1L6I were mounted on the MmGP model. The enzyme/ligand complex was prepared and energetically minimized in CHIMERA v1.19 software [28], adding the hydrogen atoms, ff14SB charges to the standard residue atoms and Gasteiger force field for nonstandard residues. Energetic minimization was performed using 1000 steepest descent steps, 10 conjugate gradient steps, both with a 0.02 Å step size and an update interval of 10.

## 3. Results

### Analysis of glycogen phosphorylase sequences

Sequence characteristics of glycogen phosphorylases from the three domains of life were analyzed by performing a protein blast. A multiple sequence alignment (MSA) was obtained for 94 sequences, and a phylogenetic tree (**Figure 1 and Figure S1**) was inferred by the maximum likelihood method. A summarized MSA, with representative sequences of Crenarchaea (*Sacharolobus solfataricus*), *Methanococcales* (*Methanococcus maripaludis* strain S2), Bacteria (*Escherichia coli*), and Eukarya (rabbit, *Oryctolagus cuniculus*) is shown in **Figure S2**. The sequences from archaea are significantly shorter than sequences from bacteria and eukarya, which results in multiple gaps with significant differences in identity percentages and sequence length across different phylogenetic groups (**Table S1**). The most striking gap in archaeal sequences is the absence of the first 150 N-terminal residues, which are reportedly involved in the dimerization and binding of allosteric effectors in other GPs [8]. Eukarya and bacteria present the well-known “large-sequence” phosphorylases of nearly 800 amino acids. In archaea, a wide range of sequence lengths are found, going from the “short-chain” phosphorylases of *Thermoproteota* with 420 residues approx. [29] to sequences of more than 800 residues in *Thermococcales* **(Table S1).**

**Figure 1.**
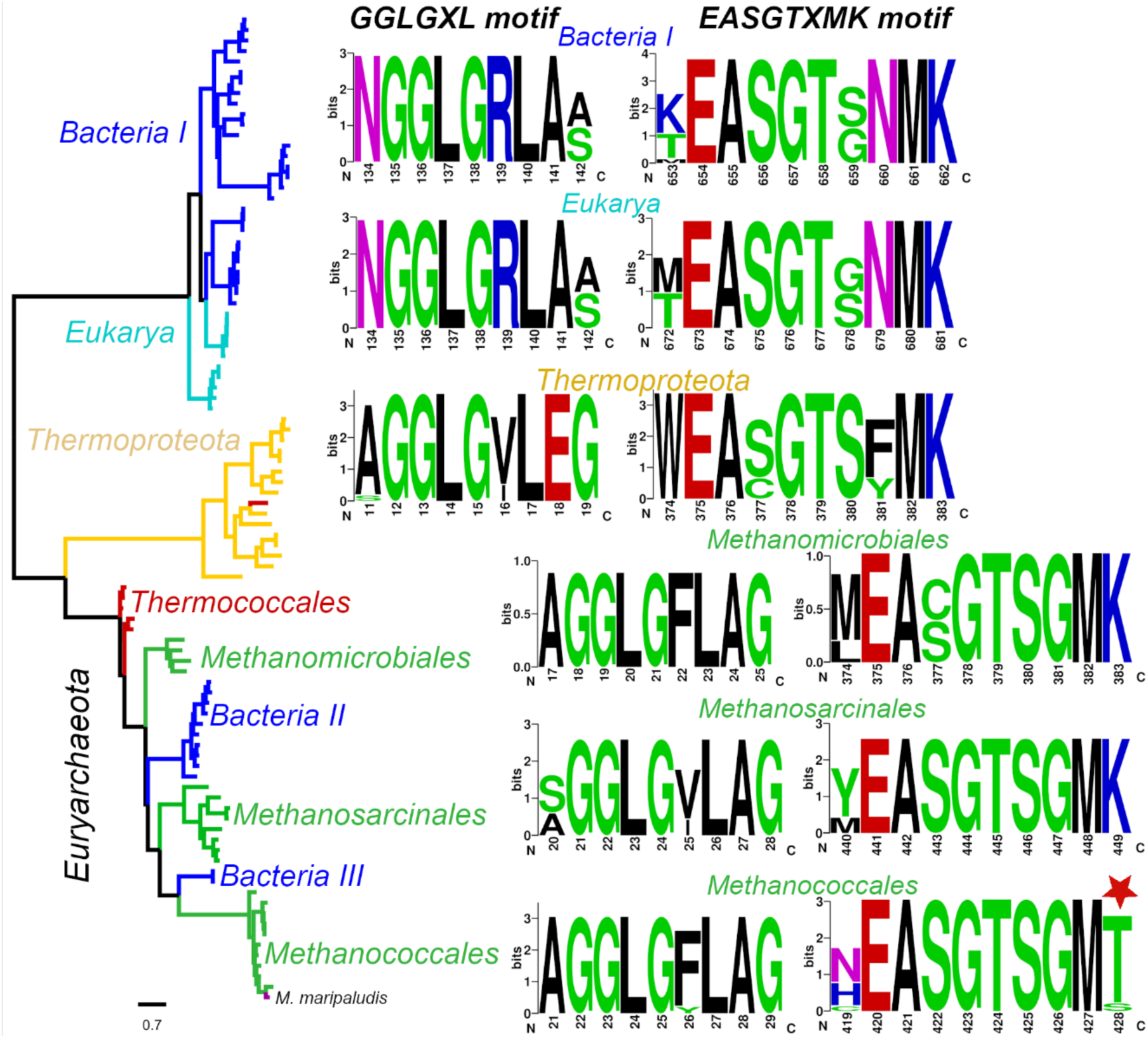
Evolutionary relationships of glucan phosphorylases and conservation of active site motifs. Phylogenetic tree for representative glucan phosphorylases from eukarya, bacteria, and archaea. SeqLogo of the active site GGLGXL and EASGTXMK motifs involved in substrate and PLP binding, respectively. Phylogeny was inferred from a sequence alignment of 94 sequences by the maximum likelihood method.

The topology of the phylogenetic tree of GP (**Figure 1**) allows the identification of three major groups of phosphorylases: (1) a group of Bacteria/Eukarya, (2) a group of only *Thermoproteota* of archaea, and (3), a group of sequences from *Euryarchaeota* archaea and bacteria (class *Actinomycetia*). The branch length between these groups indicates a large evolutionary distance, mainly between group 1 and the other groups. It is important to note that most of the experimental information available about glycogen phosphorylases comes from group 1, indicating that such information is considerably limited given the phylogenetic distance and sequence diversity we have for the other 2 groups. When analyzing the active sites, the different phylogenetic groups exhibit a high degree of conservation of active site residues despite sequence length differences, as is evident in the GGLGXL motif, involved in substrate binding and PLP positioning, and in the EASGTXMK motif (**Figure 1**), which houses the lysine responsible for covalent binding of PLP via the Schiff base [30,31]. Surprisingly, the analysis reveals that enzymes from the archaeal methanogenic order *Methanococcales* present a threonine (in some cases serine) instead of the strictly conserved lysine residue, raising questions about the functionality of GP from *Methanococcales* and their ability to bind PLP.

Biochemical and kinetic characterization of GP from *M. maripaludis*, an enzyme from the order *Methanococcales,* To determine the functionality of *Methanococcales* GPs lacking the conserved lysine active site residue, the enzyme from *Methanococcus maripaludis* (MmGP) was recombinantly expressed and characterized. The protein was purified to homogeneity according to SDS-PAGE (∼60 kDa) (Figure 2A **inset)**, and its elution volume in size-exclusion chromatography was consistent with a monomer (∼55 kDa) (Figure 2B). This differs from the oligomeric state of eukaryotic and bacterial enzymes, typically reported as homodimers and homotetramers [3].

**Figure 2.**
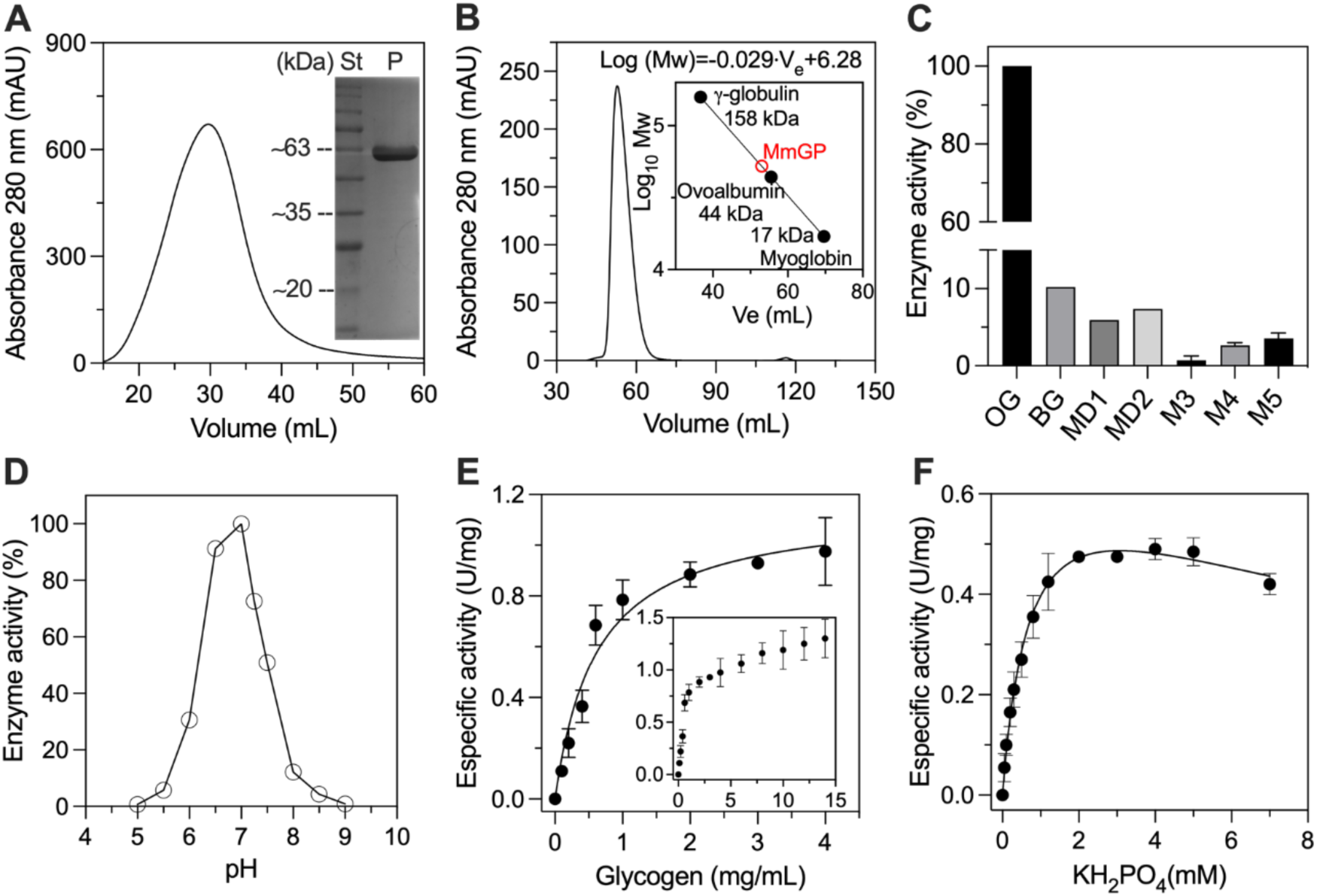
Purification and kinetic characterization of glycogen phosphorylase from *M. maripaludis*. **A)** Protein elution profile of MmGP by nickel ion affinity chromatography. Inset, SDS-PAGE of MmGP after purification (St: Molecular weight standards, P: pooled fractions). **B)** Size exclusion chromatography (Superdex S200). Inset, MW calibration curve. MmGP is shown in red (∼55kDa). **C)** Substrate preference for glucose polymers of different lengths. Oyster glycogen (OG), bovine glycogen BG, maltodextrin dextrose equivalent 4-7 unit (MD1), maltodextrin dextrose equivalent 8-15 (MD2), maltotriose (M3), maltotetraose (M4) and maltopentaose (M5). All polymers were at a concentration of 1.5 mg/mL except for maltotriose, maltotetraose, and maltopentaose which were at 3.0 mg/mL. **D)** Effect of pH on enzyme activity using a universal buffer. Substrates saturation curves at pH 7.0 for **E)** oyster glycogen using 3.5 mM of potassium phosphate as co-substrate. **F)** potassium phosphate saturation curve using 3.0 mg/mL of oyster glycogen as co-substrate.

Interestingly, MmGP showed glucose-1P production by glycogen phosphorolysis, and substrate preference analysis indicated that the highest activity was achieved with oyster glycogen, followed by bovine liver glycogen, maltodextrins (5-10% of maximal activity) and short oligosaccharides (0,7 – 3,5% of maximal activity) (Figure 2C). Structurally, oyster glycogen is more ramified than bovine liver glycogen [32,33], indicating a preference for highly branched substrates, consistent with the high polymerization reported for glycogen from prokaryotes [32]. In agreement with values reported for GP from other organisms, the pH optimum of MmGP is around 7.0 (Figure 2D) [25,34].

The kinetic parameters for MmGP were determined using oyster glycogen, and the saturation curve for phosphate shows substrate inhibition with a K_M_ value of 0.74 ± 0.13 mM. On the other hand, the glycogen saturation curve displays a Michaelis-Menten behavior up to a 4.0 mg/mL concentration, with a K_M_ value of 0.62 ± 0.11 mg/mL. At higher glycogen concentrations, the behavior deviates from the hyperbolic function and is characterized by a continuous increase in activity with no evidence of saturation (Figure 2E **inset).** This behavior might be related to the documented activation of GP by glycogen observed in the GP from rabbit muscle [35].

### Determination of pyridoxal phosphate in MmGP

Spectroscopic analyses were performed to assess if substituting the strictly conserved lysine with threonine in MmGP interferes with PLP cofactor binding at the active site. Analysis of the UV-VIS spectra of the well-characterized glycogen phosphorylase from rabbit muscle (*O. cuniculus*, OcGP) shows the characteristic absorption peak at 330 nm, which is absent in the MmGP UV-VIS spectra (Figure 3A). The fluorescence emission peak at 530, typical of PLP-containing enzymes, was also used to determine the presence of PLP. Fluorescence excitation spectra of OcGP and MmGP (measuring emission at 530 nm) show that OcGP has two excitation peaks around 290 and 335 nm and a minor peak around 425 nm, while these peaks were absent in MmGP.

**Figure 3.**
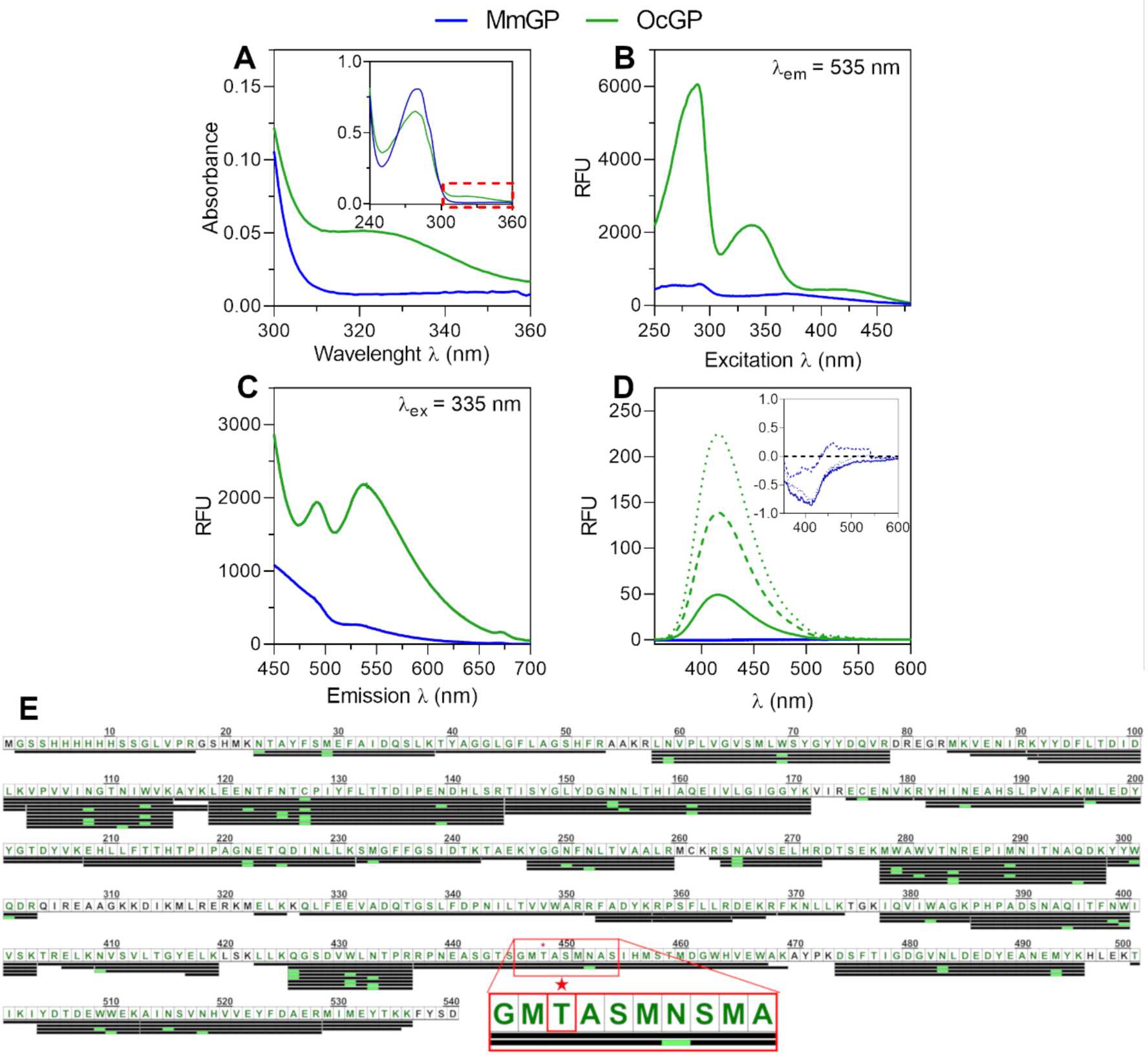
Determination of Pyridoxal phosphate content by UV-VIS, fluorescence spectroscopy, and mass spectrometry. Rabbit muscle glycogen phosphorylase (OcGP) and MmGP are denoted in green and blue, respectively. **A)** UV-Vis absorption spectra at 1 µM protein concentration for OcGP and MmGP. Inset, extended UV-Vis absorption spectra showing the absorbance peak at 280nm. **B)** Fluorescence excitation spectra recorded at a fixed emission wavelength of 535 nm at 10 µM of protein concentration. RFU: relative fluorescence units. **C)** Fluorescence emission spectra for OcGP and MmGP. **D)** Fluorescence emission spectra of the cyanohydrin adduct assayed at different protein amounts: solid line 0.2 nmol, dashed line 0.6 nmol, and dotted line 1.0 nmol. **E**) Peptide mapping by mass spectrometry of MmGP tryptic digest. The red rectangle denotes a zoom of the sequence motif that contains the lysine to threonine substitution.

Analysis of the fluorescence emission upon excitation at these three wavelengths (290, 335, and 425nm) also supports the absence of PLP in MmGP. That is, upon excitation at 290 nm of OcGP two emission peaks were observed: a main peak at 335 nm mainly contributed by aromatic residues (mainly tryptophan) and another minor peak around 530 nm characteristic of the PLP cofactor (**Figure S3A**). Excitation at 335 nm of OcGP shows two peaks around 480 and 530 nm (Figure 3C), whereas excitation at 425 nm shows the corresponding emission peak at 530 nm **(Figure S3B**). However, excitation at 290 nm of MmGP revealed only the presence of the 335 nm peak corresponding to aromatic residues (**Figure S3A**), while no emission peaks with excitation at 335 or 425 nm (Figure 3C **and Figure S3B)** were observed. Moreover, PLP content was analyzed by cyanohydrin adduct formation upon the reaction of the protein with KCN under acidic conditions [24], which allows the quantification of sub-nanomolar concentrations of PLP by following fluorescence emission at 420 nm. Analysis of the PLP content in OcGP and MmGP (between 0.2 to 1.0 nmol of protein) shows that the fluorescence of cyanohydrin adducts of OcGP correlated with protein concentration, giving a ratio of 0.83 moles of PLP per mole of protein. However, in the case of MmGP, no fluorescence emission at 420 nm was detected even with 1.0 nmol of protein (Figure 3D).

To further confirm the absence of pyridoxal 5’-phosphate or any other modification within the active site of MmGP that might explain its activity without PLP, whole-protein mass (**Figure S4**) and peptide mapping analysis (Figure 3E) was performed by mass spectrometry. The analysis reveals an observed molecular weight of 62,222 Da, roughly 178 Da smaller than the expected size of the entire protein based on the genetic sequence (62,400 Da), indicating the absence of PLP and suggesting a likely truncation or loss of one or two residues, likely at the N- or C-terminal ends. Peptide mapping (Figure 3E) achieved 89.07% sequence coverage and did not detect PLP or any post-translational modifications, neither at the active or in any other protein region. Overall, this peptide mapping analysis confirmed that the lysine-to-threonine substitution in the EASGTXMK motif of MmGP prevents PLP binding, without compromising enzyme activity.

### Proposal for the Catalytic Mechanism of MmGP

To seek a possible explanation of the catalytic activity of MmGP in the absence of PLP, we performed molecular modeling of the MmGP active site based on Alphafold2 structural prediction and comparison with the PLP-dependent maltodextrin phosphorylase of *E. coli* (EcMalP) (Figure 4), whose structure was determined with phosphate, maltopentose (M5) and PLP at the active site (PDB 1L8I). The superposition shows that in general the structure of the two proteins is quite different, with EcMalP being significantly larger than MmGP and with many non-equivalent secondary structure elements, resulting in an overall root means square deviation (RMSD) of 6.5 A. However, the structural elements of the active site are highly conserved, with the binding sites for the M5 and the phosphate substrate well conserved. On the other side, the binding site of the pyridoxal moiety of PLP is not fully conserved, as it is occupied by bulky residues such as phenylalanine 26 (F26). Interestingly, the 5’-phosphoryl group of PLP could establish polar interactions, as the surrounding residues are well conserved. To analyze the possible interactions that a phosphate group could generate at the binding site of the phosphoryl group of PLP, we mounted the phosphate group of PLP, the phosphate substrate, M5, and crystallographic waters from the active site of 1L8I to MmGP. We conducted an energy minimization without any restrictions, allowing side chains and ligands to adjust to the lowest potential energy. As its observed in the model (Figure 4), the two phosphates, M5 and several crystallographic waters generate an intricate network of polar interactions with the side chains of the MmGP active site, suggesting that could bind two phosphates at the active site, one acting as the catalytic cofactor and the other as the substrate for phosphorolysis. The two phosphates and M5 in the energetically reduced model retain their structural locations from EcMalP, as seen in Figure 4. This implies that MmGP exhibits a variant of the well-known phosphorylase [6] catalytic mechanism independent of PLP’s covalent binding.

**Figure 4.**
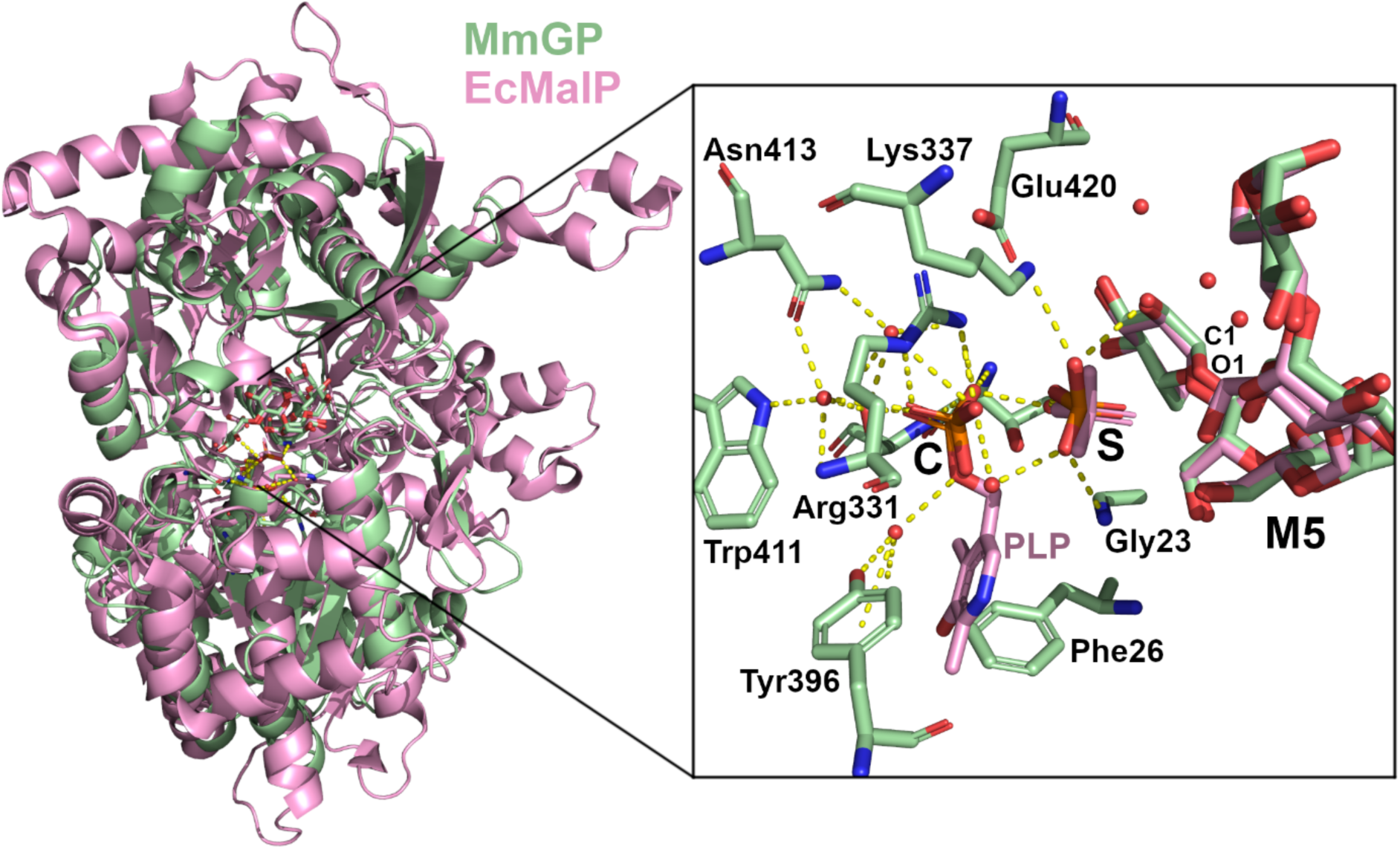
Molecular modeling of MmGP and comparison with a PLP-dependent phosphorylase (EcMalP). Structural superposition of energy minimized model of MmGP (green) and crystal structure of EcMalP (PDB 1L6I, pink). The zoom shows the active site of MmGP modeled with two phosphates: the putative catalytic one (C) and the substrate (S), in addition to maltopentose (M5). PLP, phosphate and M5 of EcMalP are shown in pink. Waters are show as red spheres.

The catalytic phosphate (C) could donate a hydrogen atom (H) to the substrate phosphate (S), and the phosphate S would donate a hydrogen atom to the O1 of the glycosidic bond. Then, the phosphate S would nucleophilically attack the C1 carbon of the first glucose residue of maltopentose. Finally, the phosphoryl group of glucose-1-Phosphate would reverse the H atom to the catalytic phosphate to complete the cycle.

### Regulation of MmGP enzyme activity

Although phosphorylases from higher eukaryotes are an archetypal model for the study of enzyme regulation, little is known about prokaryotic enzymes, which are considered as less regulated [36]. Regarding archaeal enzymes, only two have been kinetically characterized: maltodextrin phosphorylase from *T. litoralis* [37] and glycogen phosphorylase from *S. solfataricus* [29], but in none of these cases, the regulation was assessed.

To address the regulatory properties of MmGP, a set of compounds at a fixed concentration of 2 mM was assessed, including glycolytic and gluconeogenic intermediates (FBP, PEP, and PYR), glycogen metabolism intermediates (G1P, ADPG, and UDPG), Krebs cycle intermediates (citrate, malonate, and malate), nucleotides (AMP, ADP, ATP, CMP, and GMP), different phosphate forms (NaPPi and polyphosphate), and a second messenger present in *Methanococcales* (cAMP) (Figure 5A). Surprisingly, most of the assessed compounds produced changes in the enzymatic activity. All the nucleotides assayed produced a decrease in the activity, with the most significant effect produced by ADP and GMP, which decreased activity by approximately 90% and 50%, respectively, while ATP only produced a reduction of approximately 30%.

**Figure 5.**
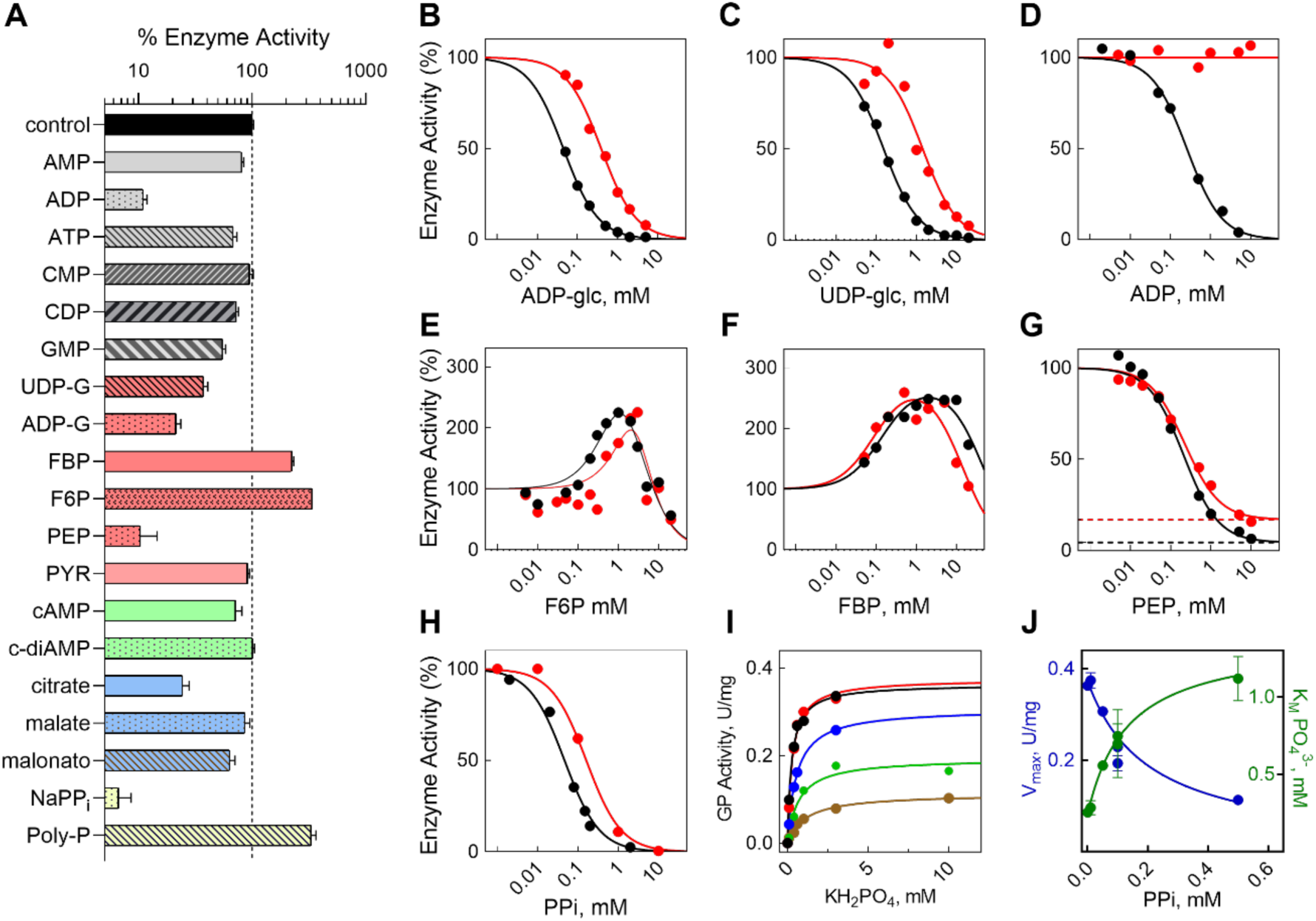
Identification and characterization of novel effectors of the MmGP phosphorylase. **A)** Screening of different molecules at 2 mM each as potential effectors of phosphorylase activity. Measurements normalized against the activity in the absence of effectors. **B-H)** Determination of the IC50 and/or AC50 for the main effectors. The curves were obtained at two substrate concentration conditions: K_M_ concentration of both substrates (0.3 mg/mL glycogen and 0.3 mM phosphate (●)) and K_M_ concentration of glycogen and saturating concentration of phosphate (0.3 mg/mL glycogen and 5 mM phosphate (●)). Equation 3 was fitted to the data in B-D and H, equation 5 was fitted to the data in E and F, and Equation 4 was fitted to the data in G. Activity values were normalized by the values of V_0_ calculated from the fitted equations. **I)** Saturation curves for Pi at different PPi concentrations (● 0, ● 0.01, ● 0.05, ● 0.1, ● 0.5 mM) at 0.3 mg/mL glycogen. The Michaelis-Menten equation was fitted, and the V_M_ and K_M_ parameters were estimated. **J)** Observed parameters V_M_ (●) and K_M_ (●) as a function of different PPi concentrations. The best fit for both parameters was a hyperbola intersecting the Y axis, corresponding to a mixed inhibition.

Regarding intermediates from glycogen metabolism, ADP-glucose, and UDP-glucose produced an important diminution in the enzyme activity, with decreases of approximately 79% and 63%, respectively. In the case of glycolytic/gluconeogenic intermediaries, PEP decreased enzyme activity by approximately 90%, while no significative effect was observed for pyruvate (PYR). On the other hand, FBP and F6P produce an increase in enzyme activity of around 338% and 915%, respectively. On the contrary, in the case of Krebs cycle intermediates, citrate decreases the enzyme activity by approximately 70%, while malate and malonate produce almost no effect. Regarding the different phosphate forms, NaPPi produces a decrease of more than 90% of the enzyme activity, but a contrary effect was observed for polyphosphate, with an increase of more than 332%. Finally, the second messenger, cAMP, produces a slight effect with only a 28% decrease in enzyme activity. These results indicate that various compounds from different metabolic pathways modify MmGP activity.

To determine the magnitude of the regulator effect, effector concentration curves were performed for the compounds showing the greatest effect (Figure 5B-H), and the IC50 (inhibitors) and AC50 (activators) values were determined **(Table S2)**. Effector concentration curves were performed at K_M_ glycogen and phosphate concentration and also at saturating phosphate concentrations (glycogen concentration near K_M_). The dose-response curves showed a total inhibition effect for ADP-glucose, UDP-glucose, ADP, and PPi. The IC50 values for ADP-glucose and UDP-glucose were 0.045 mM and 0.19 mM, respectively, and these values increased approximately 10 times at a saturating phosphate concentration. An IC_50_ of 0.24 mM for ADP was observed and the inhibitory effect disappears at saturating phosphate concentration, displaying an effect of ADP over phosphate affinity. For PPi, an IC_50_ of 0.046 mM was observed, slightly increasing 3 times when phosphate was at a saturating concentration. Noticeably, F6P and FBP behave as dual effectors, activators at low concentrations (0,1 and 1 mM), and inhibitors at higher concentrations (over 10 mM). In the case of FBP, the AC_50_ is mostly non-affected by changes in phosphate concentration, while for F6P, an increase of AC_50_ of approximately 5 times was observed. Unlike the other inhibitors, PEP behaves as a partial inhibitor (96% at K_M_ concentration and 87 % at saturation of phosphate) with IC_50_ value of 0.17 mM and presented a slight increase when phosphate concentration was at a saturating concentration.

To get insights into the mode of action of some of these effectors, we determined the inhibition mechanism of PPi, since it is highly likely to be competitive, considering its structural similarity with the putative phosphate-phosphate interaction at the active site (Figure 5I **and 5J**). Phosphate saturation curves were obtained at different PPi concentrations (Figure 5I), and the V_M_ and K_M_ values were determined by fitting the Michaelis-Menten equation. The dependence between these parameters and PPi concentration reveals a hyperbolic decrease in V_M_ and a hyperbolic increase in K_M_ (Figure 5J), showing that PPi acts as a mixed inhibitor with respect to Pi. This mixed inhibition mechanism for PPi suggests the presence of a binding site (allosteric) other than the active site for this effector.

## 4. Discussion

Glycogen phosphorylase (GP) is a key enzyme for glycogen mobilization (glycogenolysis), which have been purified from different animal, plant, and microbial sources contain PLP in a near 1:1 stoichiometric ratio [6] and the binding of PLP have been characterized by distinctive absorbance and fluorescence spectra [4,38], features that were not detected in the spectroscopic studies performed for MmGP, indicating that PLP is not present in the archaeal enzyme. This conclusion was verified by independent experiments involving PLP quantification via cyanohydrin reaction and mass spectrometry (MS) analyses. It is worth noting that previous characterization of archaeal enzymes reported the presence of PLP in enzymes from the order *Thermococcales* [37] and *Sulfolobales* [29]. The discovery that the conserved lysine of the EASGTXMK motif involved in PLP binding has been replaced by threonine in *Methanococcales* GP enzymes is surprising given the critical catalytic and conformational role of this cofactor in all phosphorylases studied thus far.

Moreover, MS analysis not only confirmed the absence of PLP but also highlighted the lack of post-translational modifications at the active site that might explain its activity without PLP, such as the introduction of a phosphate group through phosphorylation of a residue that could substitute the 5’-phosphate group of pyridoxal phosphate in catalysis.

In most PLP-dependent enzymes (i.e. transaminases, carboxylases) [4], the aldimine bond of PLP with the ε-amino group of the lysine residue of the enzyme is necessary for catalytic activity. Interestingly, the replacement of the conserved lysine has been reported for the GDP-4-keto-6-deoxymannose-3-dehydratase [39], in which a histidine replaces the lysine residue, resulting in the non-covalent binding of PLP to the enzyme. The feasibility that MmGP could bind PLP non-covalently is not consistent with UV/Vis and fluorescence spectroscopy experiments, as well as the modeling would be consistent since the pyridoxal binding site would be occupied by bulky side chains such as Phe26. Although in all phosphorylases reported, PLP also is bound by an aldimine bond to an ε-amino group of Lys [4,5], the existence of this bond is not essential for the phosphorylase reaction. Thus, the aldehyde group of PLP, which is necessary for cofactor binding to the protein, is not involved in phosphorylase catalysis. Nowadays, according to the widely accepted mechanism [6,8,40], the 5′-phosphate group of the pyridoxal phosphate acts as an acid-base to promote the attack of the substrate phosphate on the polysaccharide substrate. Therefore, our proposal that there may be two binding sites for orthophosphate may be the most plausible given the evidence, but future experiments are needed to confirm this proposal for *Methanococcales* GPs. Catalysis of the archaeal enzyme in the absence of this cofactor challenges this well-established mechanism and opens new possibilities for the glucosyltransferase reaction. This represents an unprecedented evolutionary novelty in these enzymes and raises questions about why other GP enzymes had not evolved to perform catalysis without PLP.

In addition to the direct participation of PLP in catalysis, the cofactor also plays a conformational role, as described for rabbit muscle phosphorylase, stabilizing the enzyme’s active dimeric structure and regulating the phosphorylase’s affinity to substrates and effectors [8]. Interestingly, unlike most GPs, MmGP is a monomer under the conditions studied, which ruled out a conformational role for PLP in this enzyme. The monomeric state of MmGP represents the first example of this aggregation state in GP and might be related to the absence of PLP.

Moreover, the absence of PLP in *Methanococcales* GP enzymes could have relevant biological implications. For example, it is an isolated case occurring only in this enzyme or reflects changes in the cofactor metabolism. Analysis of the PLP metabolic pathways in the BioCyC database [41] indicates that *Methanococcales* organisms harbor the pdxS/pdxT pathway for *de novo* biosynthesis of PLP and that both genes are conserved in all the organisms from this group **(Table S3)**.

Even though GP from several organisms has been thoroughly studied, and its activity is critical to the survival of some methanogen organisms under methanol exhaustion, there is no biochemical characterization of GP archaeal enzymes. The kinetic parameters of MmGP are within the range determined for prokaryotic enzymes (V_M_ 0.9 – 11 U/mg and K_M_ phosphate 1.9 – 4,0 mM) [13,25,29,37,42]. The regulation of MmGP by at least seven metabolic intermediates differs significantly from the behavior reported for bacterial enzymes, where only a small fraction responds to ADP-glucose, UDP-glucose, and, in some cases, AMP [36]. The regulation of this methanogenic archaeal enzyme is even more complex than what has been reported for eukaryotic enzymes such as yeast glycogen phosphorylase (regulated by AMP, G6P) and fungi (regulated by AMP). Some of the molecules identified in this analysis are unique regulators in MmGP, such as PEP and FBP. At the same time, ADP-glucose, UDP-glucose, PPi, and ADP have been observed to have regulatory roles as isolated cases in eukaryotic and bacterial enzymes.

Overall, the results identified a unique archaeal enzyme that differs from the well-studied enzymes from bacteria and eukarya regarding its catalytic mechanism as well as its regulation.

### 4.1 Conclusion

Glycogen phosphorylase (GP) is a key enzyme for glycogen breakdown that requires the pyridoxal 5’-phosphate (PLP) cofactor covalently bound to a strictly conserved lysine residue at the active site. Surprisingly, the enzyme from *Methanococcales* organisms presents a lysine-to-threonine substitution, representing an evolutionary novelty. Spectroscopic methods and mass spectrometry reveal the absence of the PLP cofactor in the MmGP enzyme. Despite this, the enzyme exhibits phosphorylase activity, a preference for large glucose polymers like glycogen, similar kinetic properties compared to prokaryotic enzymes, and a complex regulation by several molecules. This finding raises questions regarding the biological implications of the absence of PLP for the reaction mechanism and highlights potential new variant of the catalytic mechanisms of phosphorylases.

## Author Contributions

FGO, NHS, GVB, VCF and VG conceived the project and designed the experiments. FGO and VCF performed the phylogenetic analysis. NHS, FGO and LHC conducted the enzyme kinetic experiments. NHS, FGO and CB executed the spectroscopic, fluorometric and cyanohydrin measurements. FGO, NHS, GVB, VCF and VG wrote the article. VG and VCF were in charge of funding acquisition, project administration and supervision. All authors reviewed and authorized the final version of the manuscript.

## Acknowledgements

This work was supported by Fondo Nacional de Desarrollo Científico y Tecnológico (Fondecyt Grant 1231263 to VG and 1221667 to VCF) from the National Research and Development Agency (ANID), Chile. FGO was supported by the National Doctoral Scholarship 21191254 awarded by ANID.

## Abbreviations

AC_50_: half-maximal Activitory concentration

ADP: Adenosine 5’-diphosphate

ADPG: Adenosine-5’-diphosphoglucose

AMP: Adenosine 5’-monophosphate

ATP: Adenosine 5’-triphosphate

BG: Bovine glycogen

CMP: Cytidine 5’-monophosphate

FBP: Fructose 1,6-bisphosphate

G1P: Glucose 1-phosphate

GMP: Guanosine monophosphate

IC_50_: half-maximal inhibitory concentration

M3: Maltotriose

M4: Maltotetraose

M5: Maltopentaose

MD1: Maltodextrin dextrose equivalent 4-7 unit

MD2: Maltodextrin dextrose equivalent 8-15 unit

MmGP: Glycogen phosphorylase from *Methanococcus maripaludis*

NaPPi: Tetrasodium pyrophosphate

OG: Oyster glycogen

OcGP: Glycogen phosphorylase from *Oryctolagus cuniculus*

PEP: Phosphoenolpyruvate

PLP: Pyridoxal-5’-phosphate

PYR: Pyruvate

UDPG: Uridine 5’-diphosphate glucose

## Supplementary Material

**Supplementary Figure 1.**
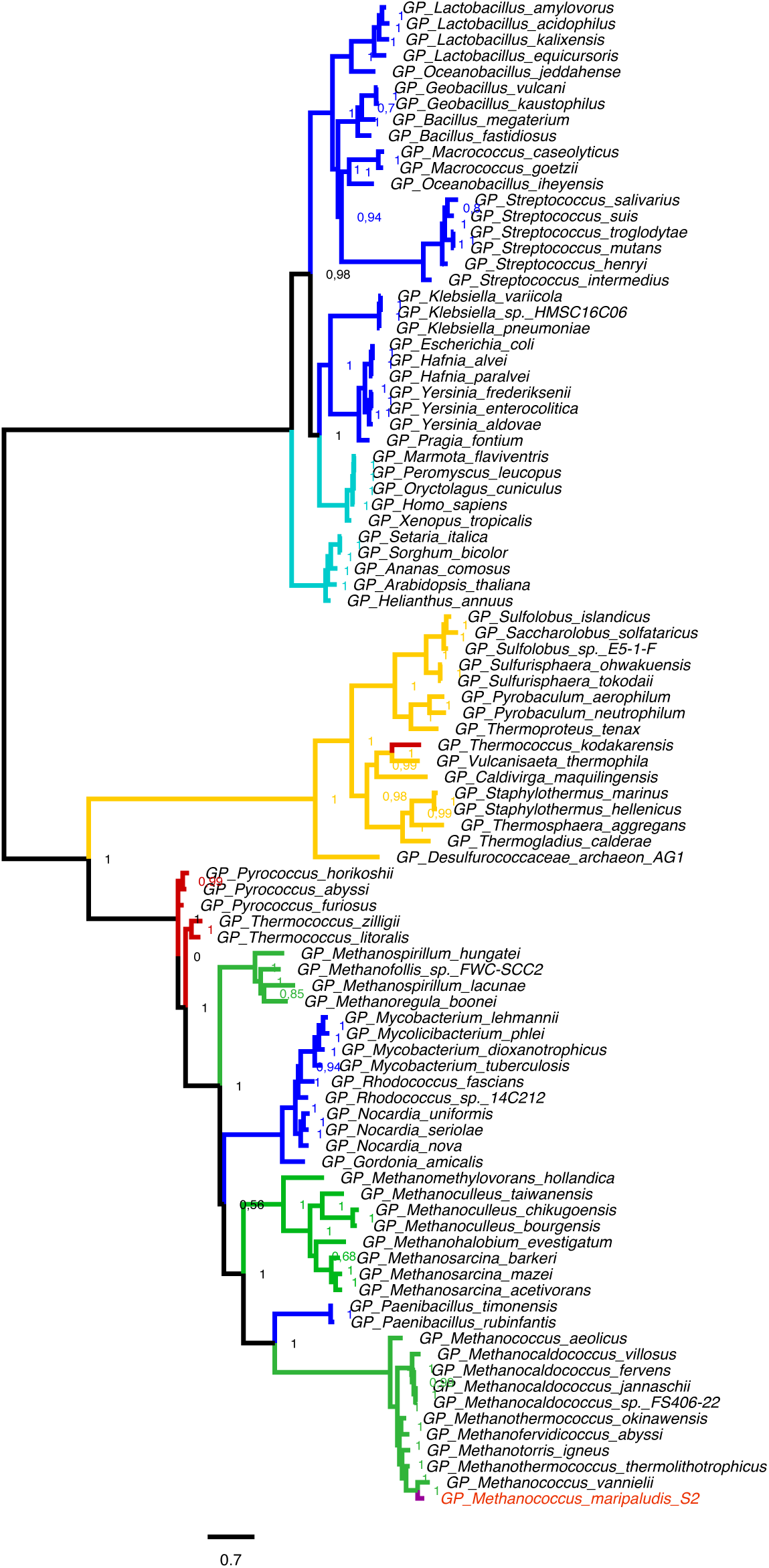
Phylogenetic tree for representative glycogen phosphorylases from archaea, eukarya and bacteria. Archaea groups are shown in different colors: Thermoproteota (yellow), Thermococcales (red) and methanogenic orders in green (Methanomicrobiales, Methanosarcinales and Methanococcales). GP sequences of the Eukarya and Bacteria domains are shown in cyan and blue, respectively. GP from M. maripaludis is highlighted.

**Supplementary Figure 2.**
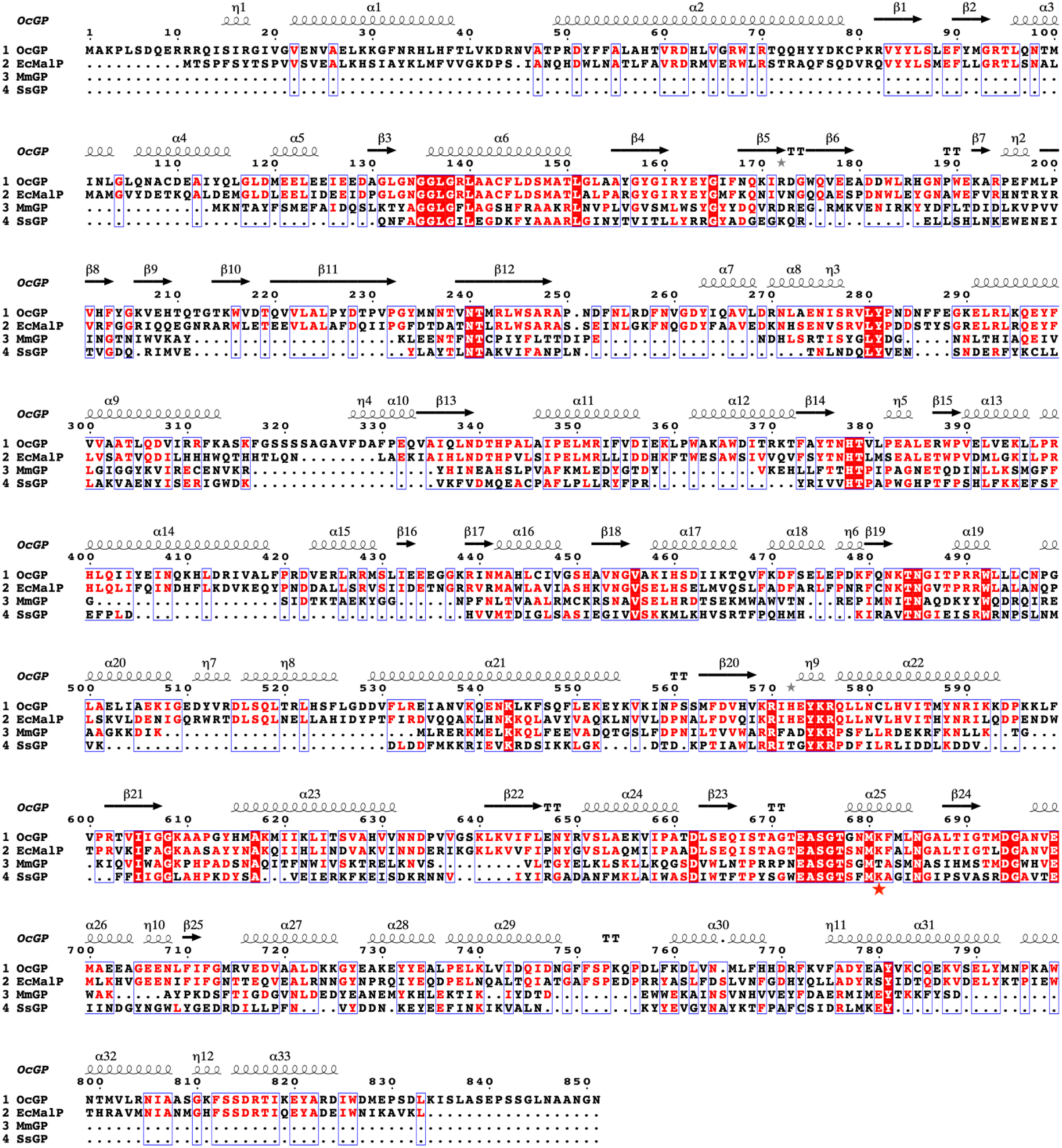
Multiple sequence alignment of representative phosphorylase sequences of eukarya, bacteria and archaea. Sequence alignment for sequences of eukarya represented by *Oryctolagus cuniculus* (OcGP), bacteria, represented by *Escherichia coli* maltodextrin phosphorylase (EcMalP), and archaea represented by *Methanococcus maripaludis* strain S2 (MmGP) and *Sacharolobus solfataricus* (SsGP). (★) Lysine to threonine replacement. The secondary structure prediction is based on the rabbit muscle phosphorylase structure PDB ID: 3ZCP.

**Supplementary Figure 3.**
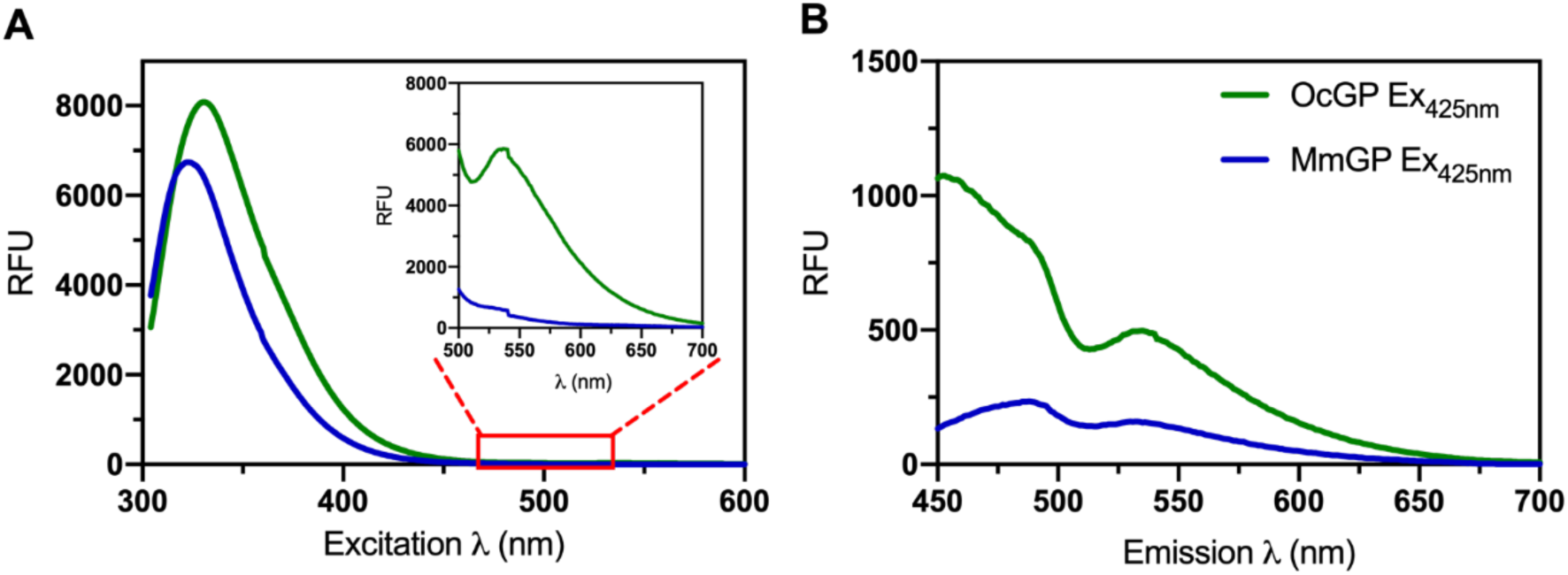
Fluorescence spectroscopy analysis of PLP content upon excitation at 290 and 420 nm. Rabbit muscle glycogen phosphorylase (OcGP) and Methanococcus maripaludis glycogen phosphorylase (MmGP) at 10 µM protein concentration, denoted in green and blue respectively. A) Fluorescence emission spectra upon excitation at 290 nm. Spectra recorded using the low sensitivity mode to compensate the high emission of the aromatic resides at 335 nm. Inset, fluorescence emission recorded in the same experimental conditions in the high sensitivity mode to enhance the emission of PLP in the 500 to 700 nm range. B) Fluorescence emission spectra upon excitation at 425 nm.

**Supplementary Figure 4.**
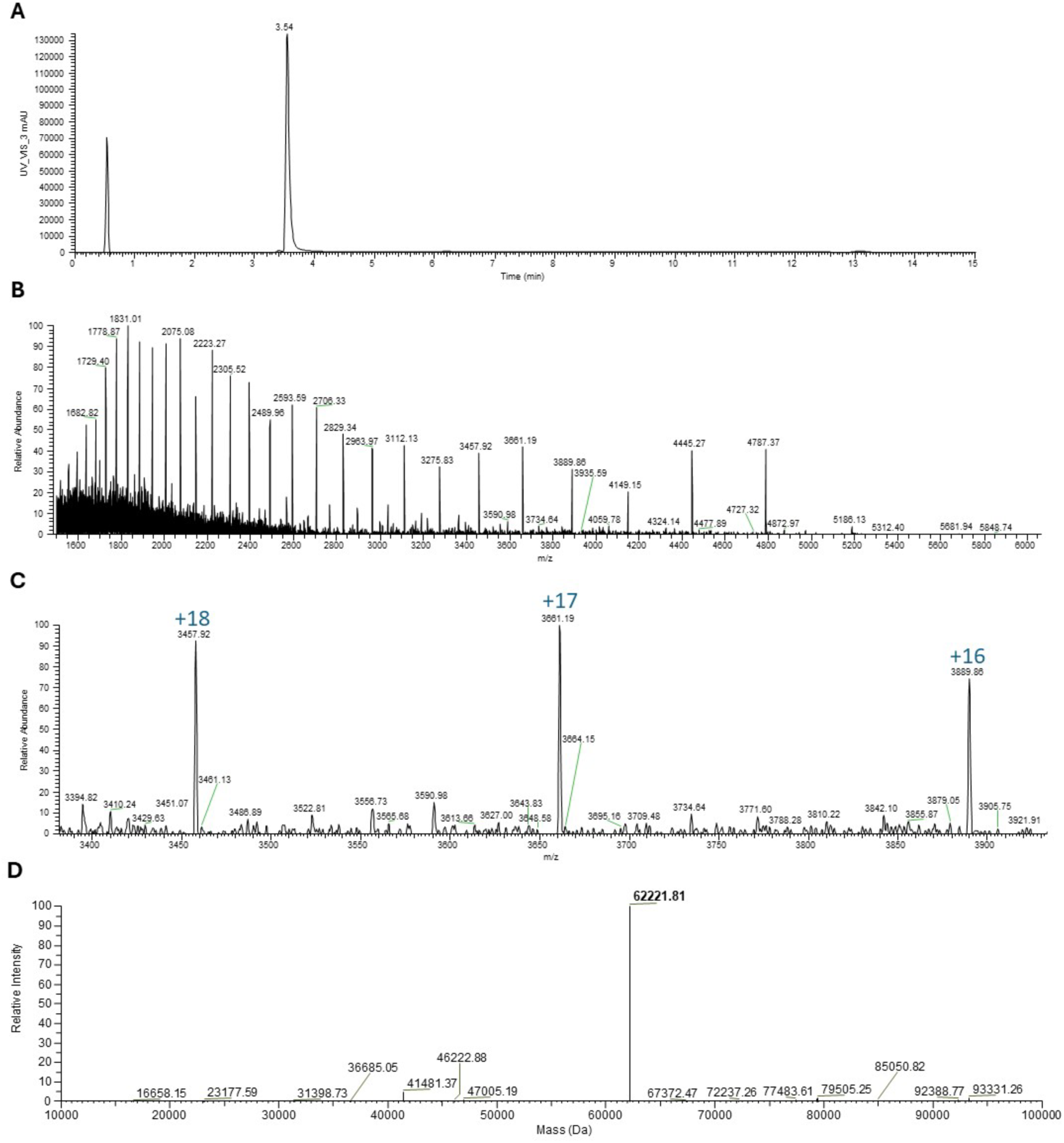
Intact mass spectrometry analysis of *M. maripaludis* glycogen phosphorylase. A) UV chromatogram (280 nm) of MmGP analyzed in a MAbPac reversed phase HPLC column. B) Coupled Q ExactiveHF-X mass spectrometry detection of mass to charge ratio. C) Zoom to the full MS spectrum. D) Mass spectrum deconvolution for the molecular weight determination of MmGP performed with the Thermo BioPharma Finder 4.0 using the ReSpect deconvolution algorithm.

---

**Supplementary Table 1.** Percentage identity and sequence length among phosphorylases from different phylogenetic groups. Percentage of amino acid sequence identity between groups was calculated between a representative sequence from each phylogenetic clade. Major differences and similarities are highlighted in red and blue respectively. The variability in the sequence length within each group is indicated by the symbol †.

|  | **Crenarchaea**<br>*S. solfataricus*<br>415 - 495 aa<sup>†</sup> | **Methanococcales**<br>*M. maripaludis*<br>518-520 aa<sup>†</sup> | **Thermococcales**<br>*T. litoralis*<br>820 – 849 aa<sup>†</sup> | **Bacteria**<br>*E. coli* MalP<br>766 -823 aa<sup>†</sup> | **Eukarya**<br>*O. cuniculus* muscle<br>760 – 861 aa<sup>†</sup> |
| --- | --- | --- | --- | --- | --- |
| **Methanococ.**<br>*M. maripaludis* | 17,93 | — | 26,76 | **11,72** | **10.92** |
| **Thermococ.**<br>*T. litoralis* | 18,54 | 26,76 |  | 13,35 | 13,76 |
| **Crenarchaea**<br>*S. solfataricus* | — | 17,93 | 18,54 | 9,16 | 9,39 |
| **Bacteria**<br>*E. coli* | 9,16 | 11,72 | 13,35 | — | 47.05 |
| **Eukarya**<br>*O. cuniculus* | 9,39 | 10.92 | 13,76 | **47.05** | — |

**Supplementary Table 2.** IC50 and AC50 values for the main effectors of MmGP idfentified IC50 values determined by fitting to a hyperbolic decay model. The error expressed corresponds to the standard deviation of the IC50 parameter. Conditions: Km/Km KH_2_PO_4_ 0.3 mM and glycogen 0.3 mg/mL; and Km phosphate/saturating glycogen KH_2_PO_4_ 0.3 mM and glycogen 2.0 mg/mL.

| Effector | IC50 (mM) |  | AC50 (mM) |  |
| --- | --- | --- | --- | --- |
|  | Substrate concentration |  | Substrate concentration |  |
|  | K<sub>M</sub>/K<sub>M</sub><br>KH<sub>2</sub>PO<sub>4</sub> /glycogen | saturation /K<sub>M</sub><br>KH<sub>2</sub>PO<sub>4</sub> /glycogen | K<sub>M</sub>/K<sub>M</sub><br>KH<sub>2</sub>PO<sub>4</sub> /glycogen | saturation /K<sub>M</sub><br>KH<sub>2</sub>PO<sub>4</sub> /glycogen |
| NaPP<sub>i</sub> | 0.047 ± 0.004 | 0.16 ± 0.02 |  |  |
| ADP | 0.26 ± 0.03 | <sup>a</sup> |  |  |
| PEP | 0.21 ± 0.03 | 0.22 ± 0.03 |  |  |
| ADPG | 0.045 ± 0.001 | 0.39 ± 0.03 |  |  |
| UDPG | 0.151 ± 0.005 | 1.4 ± 0.3 |  |  |
| F6P | 2.0 ± 1.3 | <sup>a</sup> | 1.1 ± 0.8 | <sup>a</sup> |
| FBP | 47 ± 15 | 14 ± 4 | 0.14 ± 0.04 | 0.09 ± 0.04 |

<sup>a</sup> Could not be determined.

**Supplementary Table 3.** Conservation of the pathway (pdxS/pdxT) for the *de novo* biosynthesis of PLP in *Methanococcales*. The accession numbers correspond to NCBI GenBank database.

| **Organism** | **pdxS**<br>**Accession number** | **pdxT**<br>**Accession number** |
| --- | --- | --- |
| *Methanocaldococcus*<br>*sp. FS406-22* | WP_012981081.1 | WP_012980191.1 |
| *Methanocaldococcus*<br>*bathoardescens* | WP_048202180.1 | WP_048202403.1 |
| *Methanocaldococcus*<br>*fervens* | WP_015791179.1 | WP_015791282.1 |
| *Methanocaldococcus*<br>*infernus* | WP_013100269.1 | WP_013099469.1 |
| *Methanocaldococcus*<br>*jannaschii* | WP_010870182.1 | WP_010871185.1 |
| *Methanocaldococcus*<br>*lauensis* | WP_214399381.1 | WP_214399270.1 |
| *Methanocaldococcus*<br>*villosus* | WP_004589727.1 | WP_004591278.1 |
| *Methanocaldococcus*<br>*vulcanius* | WP_015733248.1 | WP_015733095.1 |
| *Methanococcus*<br>*aeolicus* | WP_011974024.1 | WP_011974147.1 |
| *Methanococcus*<br>*maripaludis* | WP_181504408.1 | WP_181492629.1 |
| *Methanofervidicoccus*<br>*sp. A16* | WP_148120325.1 | WP_148120275.1 |
| *Methanothermococcus*<br>*okinawensis* | WP_013866750.1 | WP_048058110.1 |
| *Methanotorris*<br>*formicicus* | WP_007044953.1 | WP_007044708.1 |
| *Methanotorris*<br>*igneus* | WP_013799190.1 | WP_013798912.1 |

